# Re-evaluating Laminar Specificity of Working Memory in Human Prefrontal Cortex

**DOI:** 10.1101/2025.01.31.635930

**Authors:** Denis Chaimow, Jonas Karolis Degutis, Daniel Haenelt, Robert Trampel, Nikolaus Weiskopf, Romy Lorenz

## Abstract

Although working memory reliably activates the dorsolateral prefrontal cortex (dlPFC), the functional contributions of its cortical layers in humans remain unclear. A seminal study reported a laminar dissociation in dlPFC—superficial layers engaged during working-memory manipulation and deeper layers during motor response execution. Like many current layer-resolved studies, that work relied on several manual and semi-manual processing steps, including the selection of regions of interest. We conducted a preregistered replication in 21 human participants using an automated and fully reproducible analysis pipeline. While we replicated the superficial-layer effect during manipulation, we found no evidence for preferential deep-layer activation during response execution. This biologically plausible result refines current models of laminar organization in the human prefrontal cortex and aligns human evidence with the more heterogeneous animal literature. By demonstrating reproducibility through preregistration and automation, this work establishes a benchmark for laminar analyses in cognitive neuroscience.

## Introduction

The dorsolateral prefrontal cortex (dlPFC) plays a central role in human high-level cognition, consistently activating during working memory (WM) tasks that require the temporary retention and manipulation of information(Curtis & D’Esposito, 2003; D’Esposito & Postle, 2015). Despite its significance, the functional role of the dlPFC’s laminar organization in human WM remains poorly understood.

Evidence from invasive electrophysiology in non-human primates suggests that distinct layers of the dlPFC contribute differently to WM processing, although findings across studies are not uniform. Delay-period activity has often been localized to superficial layers 2–3 (Bastos et al., 2018; Sawaguchi et al., 1989, 1990), giving rise to models that attribute WM delay activity to recurrent interactions among nearby, similarly tuned cortical columns (Compte et al., 2000; Goldman-Rakic, 1995; Kritzer & Goldman-Rakic, 1995). In contrast, neurons in deep layer 5 have been shown to be more active during motor responses following the delay (Opris et al., 2011). However, this pattern is not exclusive. Other studies have reported response-related activity in superficial layers (Bastos et al., 2018; Markowitz et al., 2015), indicating that both compartments can contribute across task epochs. Together, these data point to interacting rather than strictly segregated laminar contributions to WM and highlight that laminar specialization in dlPFC is complex and context-dependent.

The development and validation of such mechanistic models of cortical function depend on the ability to measure neuronal activity at the resolution of cortical layers. Until recently this has been the exclusive domain of invasive electrophysiological studies and therefore limited to non-human animals. Animal studies, while mechanistically rich, remain limited in scope: they typically rely on years-long training in narrowly constrained tasks (Yang et al., 2019), recordings from a small number of cortical sites, and species with fundamentally different cognitive repertoires. Moreover, intensive training reshapes prefrontal representations in humans (Miller et al., 2022), raising concerns about generalizing from overtrained animal models. Thus, translating these insights into a mechanistic framework for human cognition remains a critical challenge.

Advances in high-resolution functional neuroimaging have now paved the way to study the human brain at laminar resolution. Most layer fMRI studies so far have targeted sensory (Kok et al., 2016; Koopmans et al., 2010; Olman et al., 2012) and motor systems (Huber et al., 2017), but recent work has extended these methods to higher-order association cortices (Degutis et al., 2024; Finn et al., 2019; Sharoh et al., 2019). In their seminal study, Finn et al. (2019) demonstrated layer-specific responses in the human dlPFC for the first time during a WM task using high-resolution VASO fMRI. This imaging sequence offers higher spatial specificity than conventional gradient-echo BOLD, making it well suited for laminar analyses. Finn et al. reported that superficial layers were more active during WM manipulation, whereas deeper layers were more active during motor response execution—a pattern interpreted as distinct laminar computations consistent with models derived from animal work. This clear dichotomy has strongly influenced subsequent human laminar research, shaping ongoing efforts to link cortical microcircuit mechanisms to cognition.

Finn et al. (2019) used an analysis approach involving careful manual localization of active regions of interest (ROIs) and layer delineation in dlPFC. Such manual and semi-manual processing steps are common in many layer fMRI studies to date (Dresbach et al., 2023; Faes et al., 2023; Huber et al., 2023; Zhang et al., 2023), aiming to achieve high precision. This is understandable as until recently the focus has been on feasibility and demonstration studies. However, these approaches are labor-intensive, prone to subjectivity, often non-preregistered, and applied to small samples, leaving many laminar findings awaiting replication. To make the leap from methodological demonstration to routine use in cognitive neuroscience,it is essential to establish the replicability of key results using automated and reproducible pipelines.

Here, we conducted a preregistered replication of this seminal layer-specific working memory study in humans using ultra-high field VASO fMRI at 7 Tesla and a fully automated analysis pipeline. Across 21 participants, we replicated the superficial-layer effect during working memory manipulation but found no evidence for preferential deep-layer activation during motor response execution. Extensive control analyses confirmed the robustness of these results. Our findings refine current models of laminar organization in the human dlPFC, demonstrating that superficial-layer activity is a reproducible signature of working memory manipulation, whereas deep-layer effects appear less robust - a pattern consistent with the heterogeneous and task-dependent laminar profiles observed in animal studies. By combining preregistration, automation, and a fully transparent analysis workflow, our study provides a methodological benchmark for reproducibility in layer fMRI and discusses limitations inherent to this technique, while strengthening the foundation for linking human laminar organization to mechanistic models derived from animal neurophysiology.

## Results

### Preregistered replication analysis

We determined the target sample size of 21 participants using conservative power analysis based on effect sizes of the original study (see Methods). Participants performed a WM task identical to Finn et al. (2019), while we simultaneously acquired VASO and BOLD fMRI data at 7T using an SS-SIVASO sequence (Huber et al., 2014; Huber et al., 2021; Stirnberg & Stöcker, 2021). Briefly, in each trial of two experimental runs a random set of five letters was presented visually, followed by a cue indicating whether the letters had to be mentally rearranged in alphabetical order or merely remembered in their original order (alphabetization vs. remembering). After a delay of 10 s one of the letters appeared as a probe and participants had to press one of five buttons indicating its alphabetical or ordinal position. In two additional runs only alphabetization trials were presented but were varied with respect to whether a response was required or not (action vs. non-action).

We conducted a replication study of Finn et al. (2019) following an analysis pipeline that we had preregistered on Open Science Framework Registries (https://doi.org/10.17605/osf.io/txh3s). We designed the analysis pipeline to closely follow the original study. However, consistent with our overall goal, we formalized all manual steps. Most notably, ROIs were automatically determined by combining precise anatomical information about dlPFC localization obtained from a fine-grained surface-based cortical parcellation (Glasser et al., 2016) with subject-specific clusters of task-evoked functional activation in the BOLD data (Fig. 1 and Methods). Following Finn et al. (2019), we leveraged BOLD fMRI for the ROI selection procedure due to its higher sensitivity compared to VASO fMRI. We then split each automatically defined subject-specific ROI into superficial and deep layer voxels based on cortical depth estimates (Fig. 1) and estimated average trial time courses from the VASO data as a function of layer and experimental condition.

**Figure 1.**
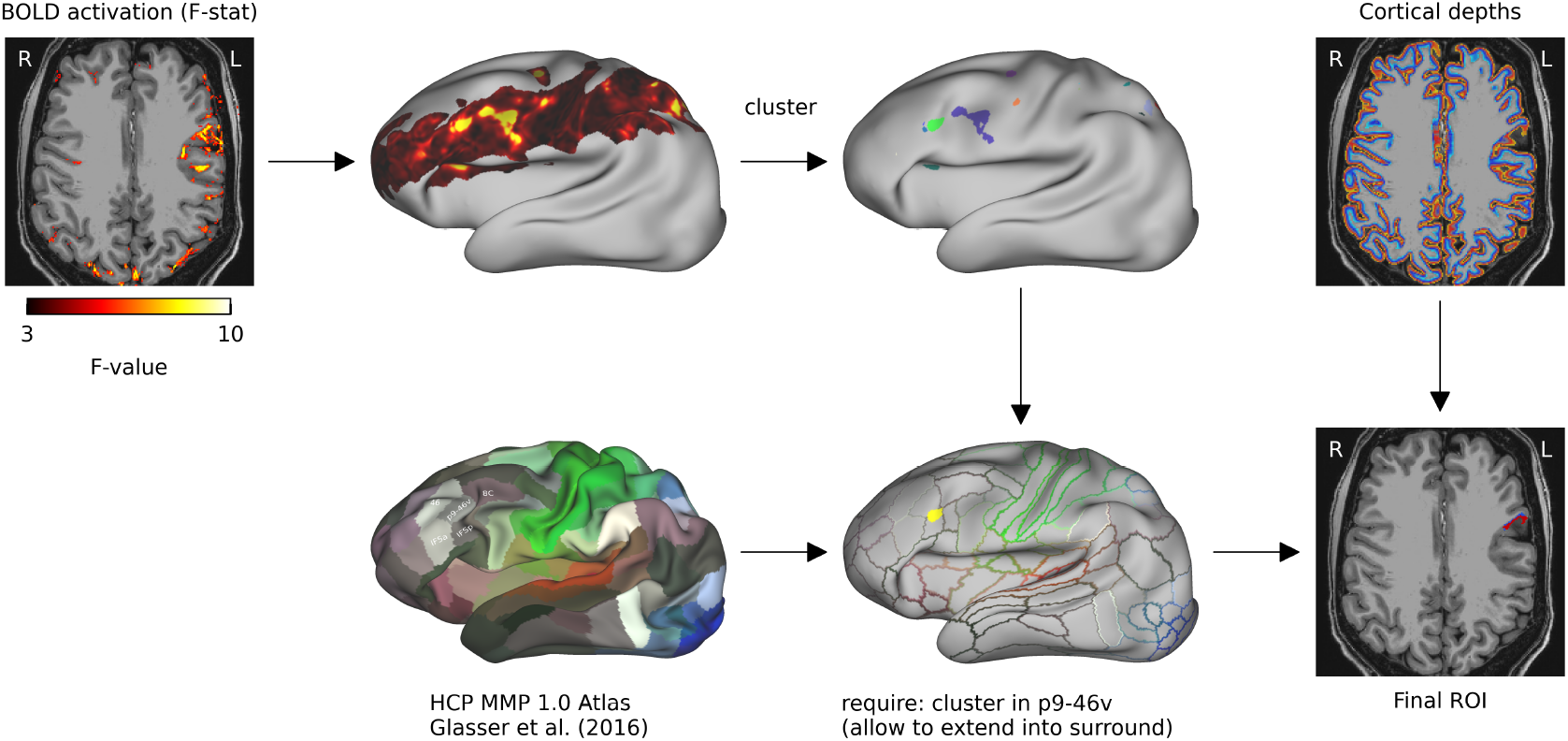
Automatic definition of ROIs. ROIs were determined by combining anatomical information obtained from the HCP MMP 1.0 atlas (Glasser et al., 2016) (bottom left) and on whether a location was activated by the experimental paradigm using the BOLD contrast (top row from left). We chose the largest activation cluster overlapping with the dlPFC, corresponding to the p9-46v parcel following Finn et al. (2019), and selected all contiguous voxels that belonged either to p9-46v or any of the surrounding parcels (46, 8C, IFSp, IFSa) accounting for uncertainties in single subject parcellation. This ROI was then split into a superficial and a deep layer based on cortical depth estimates.

We present the resulting laminar time courses averaged over subjects (Fig. 2A) in a format similar to Finn et al. (2019). The alphabetization condition resulted in elevated VASO signals following the delay period as compared to mere remembering (compare blue to green curve). Furthermore, action trials resulted in elevated VASO signals following the response period (compare red to orange curve). To quantify these differences, we first averaged the time points associated with delay and response periods (Figure 2B) while accounting for hemodynamic delay. We then analyzed the dependence on cortical layer, experimental condition, and trial period using a series of two-way ANOVAs, identical to Finn et al. (2019). First, we analyzed each layer’s response in each type of run separately for effects of experimental condition and trial period.

**Figure 2.**
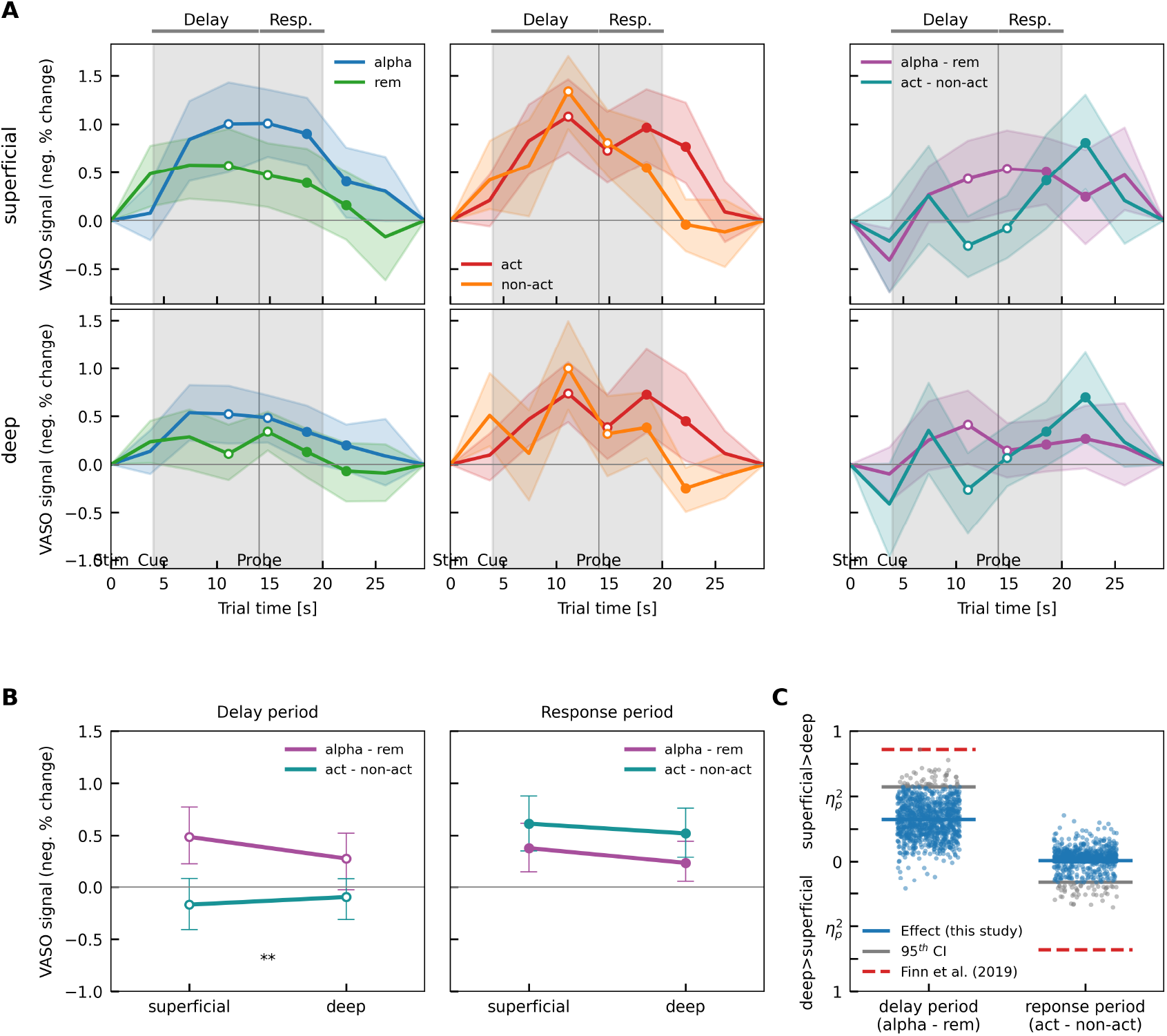
Results of preregistered replication analysis. **(A)** Trial averaged VASO time courses for superficial (top row) and deep layers (bottom row) for alphabetizations vs. remembering (left), action vs. non-action (middle) and differences between conditions from both types of runs. Each panel shows the negative percent VASO signal response, i.e. a proxy of positive cerebral blood volume change and activation (see Methods), as a function of trial time. A sequence of 5 random letters was presented at t = 0 s. At t = 4 s a cue indicated whether participants had to alphabetize the sequence or remember it in its original order. At t = 14 s one letter was probed, and participants had to respond using a button press. The shaded band indicates the 95% confidence interval over subjects. Open circles and filled circles represent delay and response period data points, respectively, that were used for further analysis **(B)** VASO response contrasts obtained from delay (left) and response periods (right) are shown for alphabetization vs. remembering (purple) and action vs. non-action (teal) for superficial and deep layers. The interaction between type of contrast and layer was significant for the delay but not for the response period. **(C)** Estimated interaction effect sizes (partial eta squared; blue lines) for delay and response periods are shown together with their sampling distributions estimated using bootstrapping (blue circles). Effect size values belonging to a higher superficial layer contrast are plotted upwards, those belonging to a higher deep layer contrast downwards. For both periods the interaction effects’ 95^th^ percentiles (gray lines) are clearly distant from the results of Finn et al. (2019), shown as dashed red lines.

### Replicating superficial layer delay effect but not deep layer action effect

To determine whether response differences between conditions were layer-specific and dependent on the type of experimental manipulation, we conducted two preregistered two-way ANOVAs, following methodology outlined by Finn et al. (2019). For each task period, we tested for an interaction between layer (superficial vs. deep) and condition contrast (alphabetization vs. remembering for the delay; action vs. non-action for the response). This constitutes the main analysis targeted by our preregistered power calculation We found a significant interaction for the delay period (p=5.8×10^−3^; F(1,20)=9.53; η_p_ ^2^=0.323) but not the response period (p=0.70; F(1,20)=1.52×10^−1^; η_p_ ^2^=7.57×10^−3^). As in Finn et al. (2019), the interaction pattern during the delay reflected stronger superficial-layer responses for the alphabetization compared to the remembering condition. However, contrary to the original study, no preferential deep-layer activity was observed during the response period; instead, superficial layers showed a weak trend toward higher activation for action versus non-action trials.

Supplementary analyses examined main effects of condition and period within each layer and confirmed higher overall activity for alphabetization compared to remembering trials across both layers, as well as action-dependent increases in superficial layers during the response period (see Supplementary Results 1 and 2 for detailed statistics).

Though not part of the preregistered replication analysis, we carried out the same statistical analyses on the BOLD part of the data (Supplementary Fig. 1). Here, significant interaction effects were observed, with both alphabetization and action conditions, both showing stronger responses in superficial layers (delay period: p=9×10^−6^, F(1,20)=34.6; η_p_ ^2^=0.634; response period: p=2.9×10^−3^, F(1,20)=11.5; η_p_ ^2^=0.364; two-way ANOVA interaction with factors layer and condition contrast). However, these results pointing to relative higher activity in superficial layers need to be interpreted with caution as BOLD is known to be affected by draining-vein bias leading to an overrepresentation of superficial layer activity (Degutis et al., 2025; Havlicek & Uludağ, 2020; Heinzle et al., 2016; Markuerkiaga et al., 2016).

### Failure to replicate deep layer action effect not due to limits in sensitivity

A lack of a statistical interaction effect for the response period in the VASO analysis might suggest that any true effect present in our data was smaller than previously reported, or that our study lacked the statistical power to detect effects comparable to those reported by Finn et al. (2019). To further compare our results to Finn et al. (2019) we used bootstrapping over subjects to estimate confidence intervals on the layer by condition contrast interaction effects, taking the direction of layer differences into account. For the superficial layer delay effect, 95% of bootstrap samples yielded η_p_^2^ values below 0.574 or failed to demonstrate an increased superficial layer delay contrast (blue dots below the zero line in Fig. 2C). These findings stand in contrast to the value of 0.86 reported by Finn et al. (2019). For the deep layer action effect, 95% of bootstrap samples either did not result in any higher deep layer action contrast or yielded η_p_^2^ values below 0.164, compared to an effect size of η_p_^2^=0.68 reported by Finn et al. (2019) (Fig. 2C). This discrepancy suggests that any potential deep layer action effect is considerably smaller than reported by Finn et al. (2019) and that their superficial layer delay effect is also stronger than what our data suggests.

### Control analyses demonstrate adequacy of methods and robustness of results

We conducted a series of control analyses to evaluate how data quality and methodological choices may have influenced our results. First, we assessed our data quality by calculating temporal signal-to-noise ratio (tSNR) maps on individual VASO runs (Supplementary Fig. 2) and averaging the within-ROI tSNR for each subject. The median tSNR across subjects was 14.1, which can be considered moderate to excellent (Huber, 2020).

We also inspected gray matter segmentation and structural to functional image registration focusing on each participant’s individual ROI (Supplementary Fig. 3). The fully automatic segmentation and registration appears to have performed well within and around each ROI. However, in subjects 2, 3, 4 and 7 the automatically generated ROI was located near the edge of the slab, resulting in lower tSNR (Supplementary Fig. 2) and reduced ROI volume (Supplementary Fig. 6a). We therefore reran our main analysis without these four subjects. Results for this subgroup were similar to those from the complete group (Supplementary Fig. 4) in that there was a significant interaction for condition contrast and layer during the delay period (p=0.011; F(1,16)=8.38; η_p_^2^=0.344) with the superficial layer showing higher responses specific to the alphabetization vs. remembering contrast. As in the main analysis there was no significant interaction during the response period (p=0.87; F(1,16)=2.5×10^−2^; η_p_^2^=1.61×10^−3^).

We have thus qualitatively demonstrated that the automatic segmentation and ROI definition performed reasonably well. To further quantitatively assess potential loss of accuracy compared to manual segmentations we also created manually drawn laminar ROIs. We hand-segmented superficial and deep layers directly on T1-weighted VASO images, a procedure comparable to Finn et al. (2019), but we used the automatic ROIs to determine location and extent. This procedure required about 50 hours of manual labor. We reran our main analysis using these carefully manually delineated ROIs. Resulting time courses (Supplementary Fig. 5) were qualitatively very similar. Importantly, we did not find a deep layer action effect (p=0.52, F(1,20)=0.431, η_p_ ^2^=2.11×10^−2^). Notably, the superficial layer delay effect size decreased compared to the automatic ROI analysis (p=0.09; F(1,20)=3.16; η_p_ ^2^=0.136). Although the decrease of the superficial layer effect was somewhat unexpected, the confidence intervals of the manual and automatic analyses highly overlap (Supplementary Fig. 5C). Together these results point against limited sensitivity of our automatic ROI analysis approach.

In our automated ROI estimation we had to define a target ROI surface area. It is possible that smaller ROIs would have been more sensitive to a more localized effect. Alternatively, assuming a more distributed response, averaging over a larger ROI could have boosted statistical power. Supplementary Fig. 5 shows the effect of varying ROI size. We find some variations in the strength of the superficial delay layer effect during the delay period in that moderately small surface areas between 80 mm^2^ and 120 mm^2^ yield statistically significant results while very small surface areas (< 80 mm^2^) as well as surface areas between 140 mm^2^ and 320 mm^2^ are not significant; for very large areas from 340 mm^2^ on, results would again yield statistical significance. Importantly though, no ROI size resulted in a deep layer action effect.

In our analysis, we followed the ROI description provided by Finn et al. (2019), where the focus was on activations in and around the left dlPFC (approximately Brodmann area 9/46) which we preregistered as the left p9-46v parcel from the HCP MMP 1.0 atlas (Glasser et al., 2016). However, it is important to note that their subject-specific ROIs were drawn manually and without the help of atlasing tools, and these ROIs are not publicly available. Given that these ROIs were manually defined, there is a possibility that the original analysis inadvertently concentrated on alternative dlPFC ROIs. In light of this, we should reconsider the focus on p9-46v and explore other candidate regions of the dlPFC to ensure a more comprehensive and accurate assessment of task-specific activations.

To that extent, we estimated clusters in the group averaged BOLD F-stat map, backprojected them to individual subjects and used these as constraining regions for automatic ROI calculation. Supplementary Fig. 7 shows that these clusters resemble the multiple demand (MD) system (Assem et al., 2020; Duncan, 2010), a frontoparietal network that typically activates during a wide variety of cognitively demanding tasks. Clusters MD3A and MD3B resemble our main ROI region in and around left p9-46v. In addition, clusters MD2 and MD4 situated in close posterior and anterior proximity, respectively, are candidates for alternative dlPFC ROIs. However, the ROIs based on any of these group clusters resulted in qualitatively similar layer and condition contrasts as our main ROI (Supplementary Figs. 8-11). In particular, none of them showed a deep layer action effect.

Finally, we assessed whether a larger separation of layers within the ROI would affect results by increasing laminar specificity. We reran the main analysis on modified laminar ROIs by leaving a gap of ⅓ relative cortical depth between the superficial and deep layer ROI. Results (Supplementary Fig. 12) were qualitatively similar to the main analysis. Again, superficial layers showed a higher alphabetization - remembering contrast during the delay period (contrast x layer interaction: p=1.7×10^−2^; F(1,20)=6.74; η_p_^2^=0.252). While there was a small interaction effect for the response period (p=0.50; F(1,20)=0.477; η_p_^2^=2.33×10^−2^), action-related signal increases were again higher in superficial than in deep layers.

## Discussion

In this preregistered study, we conducted a replication of a seminal study that first demonstrated layer-specific responses in the human dlPFC during WM (Finn et al., 2019). We were not able to fully replicate the results of Finn et al. (2019) using our preregistered analysis. While we replicated the findings of stronger superficial-layer activity during WM manipulation compared to maintenance during the delay period, we found no evidence for preferential deep-layer activation during motor response execution. Exhaustive control analyses all converged on this conclusion, indicating that the failure to replicate was unlikely due to low sensitivity, inaccuracies in automatic layer ROI segmentation, or differences in ROI size or definition.

Finn et al. (2019) suggested that their findings regarding superficial layer activity at least partially reflect recurrent excitatory connections. Our findings, showing a less distinct higher superficial layer response during manipulation trials, do not contradict their conclusion. However, it questions the degree to which manipulation-dependent increases in delay period activity are exclusively localized to superficial layers. Evidence from non-human primates indicates that some delay period activity is also present in deep layers, albeit to a lesser extent. Sawaguchi et al. (Sawaguchi et al., 1990) using single cell recordings during an oculomotor delayed response task found that, while most delay-type neurons that did not differentiate between stimuli were found in superficial layers, most stimulus differentiating delay period neurons were found in deep layers. Similarly, deep layer delay period activity has been found in LFP and multi-unit activity (MUA) in multiple prefrontal areas (Bastos et al., 2018). These findings also align with our previous work in humans using BOLD fMRI in WM tasks. Specifically, we observed activation in the deep layers of the left dlPFC during the delay period for heightened WM load demands, albeit to a significantly lesser extent than in the superficial layers (Degutis et al., 2024). Importantly, in our previous work, we demonstrated that this pattern is generalizable to increases in WM load more broadly and does not appear to be specific to WM manipulation, further supporting the robustness of our findings.

In contrast, our findings did not reveal stronger deep layer responses for action compared to non-action trials during the probe period. Instead, we observed increased activity in both layers but especially superficial. These results do not rule out a deep layer involvement but also do not support conclusions about an exclusive deep layer response. We speculate that such distinct deep layer responses are also not very likely. Even though there is ample evidence from electrophysiological studies in non-human primates that dlPFC deep layers are active and involved in generating a response in delayed-response tasks (Opris et al., 2011; Sawaguchi et al., 1989, 1990), findings across studies are heterogeneous and do not support a strictly layer-specific dissociation. Delayed response tasks in non-human primates also showed superficial layer activity well into the probe period (Bastos et al., 2018; supplementary Fig. 8). This was true for a search task in which images had to be compared to WM content but also for a simpler delayed saccade task in which cognitive demands after the delay were minimal. Similarly, Markowitz et al. (2015) isolated WM storage-specific population activity, that could be localized preferentially to superficial layers, and that showed heightened activity not only during the delay period but also during the probe period when a memory-guided saccade had to be made. Together, these findings indicate that superficial-layer engagement during the probe period is not uncommon in delayed-response paradigms, aligning with our finding of heightened superficial activity during action conditions.

Apparent discrepancies with reports of exclusive deep-layer responses in animal electrophysiology may also reflect differences in signal origin. Electrophysiological recordings capture spiking activity, which primarily reflect action potentials originating from the somata of the neurons. In contrast, fMRI signals are more closely related to local field potentials (Logothetis et al., 2001), which primarily reflect synaptic activity and dendritic integration, potentially extending into more superficial layers as is the case for apical dendrites of layer 5 pyramidal neurons. If the deep layers of the dlPFC are indeed responsible for motor response generation, the fMRI signal in the deep layers themselves might be weaker than expected as it primarily reflects synaptic input and dendritic activity rather than axonal output or somatic spiking, with stronger signals potentially arising in downstream targets where the motor commands are processed. However, this might not be the case for feedback connections that target deep layer synapses and could explain why layer fMRI studies have found deep layer-specific results in sensory cortices in the past (Carricarte et al., 2024; Kok et al., 2016).

In addition to these physiological considerations, the paradigm used by Finn et al. (2019) introduces another potential factor that could influence laminar activation patterns: action trials differed from non-action trials not only in the requirement of a motor response but also in whether the probed letter was shown (action) or not (non-action). This points to a potential drawback of the experimental paradigm as it complicates the clear dissociation between motor response and retrieval, making it difficult to isolate the specific contributions of each cognitive process. Previous findings have shown ramping gamma-band activity at the end of memory delays, which is thought to result from its disinhibition following a drop in alpha/beta power, allowing for a successful read-out of information from WM (Buschman & Miller, 2022; Hussar & Pasternak, 2010). Since gamma-band activity has been linked to superficial layers (Bastos et al., 2018; Buffalo et al., 2011; Maier et al., 2010), this could explain why we did not observe exclusive deep layer activation in our study. Specifically, the readout from WM during the retrieval period in the action conditions likely contributed more prominently to superficial layer activity, compared to the non-action conditions where no such readout was required. Converging evidence from a recent laminar fMRI study by our group supports this interpretation (Degutis et al., 2024). In that study, we dissociated WM retrieval from motor processes by always presenting a probe and cuing participants to either “respond” or “abstain” at retrieval. While we similarly did not observe any exclusive deep-layer effect, we also did not find heightened superficial-layer responses during motor execution, neither in overall signal strength nor in the fine-grained activity patterns using decoding analyses. In addition, we found that increased WM readout demands were associated with superficial layer processing, indicating that retrieval and comparison operations primarily recruit superficial circuits of the dlPFC.

Beyond these functional interpretations, it is important to acknowledge methodological factors inherent to laminar fMRI that may further contribute to the absence of isolated deep-layer effects. Even if laminar neuronal activity were perfectly localized to individual layers, a certain degree of laminar unspecificity in fMRI measurements is expected. Reasons are residual effects of subject motion and the sampling of cortical depths across approximately 2 mm cortical thickness into 0.8 mm or 0.9 mm wide voxels. The latter is expected to introduce some amount of cross-layer blurring (e.g. see simulations in Koopmans et al., 2011; Shmuel et al., 2007) due to varying alignment between voxels and local cortical geometry as well as biological and measurement uncertainty in exact layer boundaries. For deep layer activity there is the added effect of draining veins, whereby spatially specific vascular changes in deep layers are being propagated downstream by ascending cortical veins (Degutis et al., 2025). This effect is most significant for deoxyhemoglobin changes measured by GE-BOLD fMRI. However, some minimal sensitivity to blood volume changes in larger vessels, foremost intra-cortical arteries is expected when using VASO fMRI (Akbari et al., 2022). While varying degrees of task-dependent coactivation of deep and superficial layer activity has been demonstrated (Huber et al., 2017), to date we are not aware of any VASO results showing completely isolated deep layer responses compared to baseline, except for Finn et al. (2019).

Here, we developed a fully automated analysis while still focusing on precision. Nevertheless, it may well be the case that results like those presented in Finn et al. (2019) are currently only achievable with careful manual intervention. However, it should be noted that previous attempts of automated segmentation were able to yield layer-specific results (Degutis et al., 2024; Kok et al., 2016; Lawrence et al., 2019). Moreover, our manual delineated ROIs did not reveal qualitatively different results. We believe that manual approaches play an important role in advancing current analysis techniques and are therefore well justified as an initial step. However, the next step must be to give such methods and results a solid foundation that ensures replicability with adequate methodological effort. Defining what constitutes “adequate” requires a deeper exploration of how various methodological choices in data acquisition and analysis impact the outcomes.

General challenges in layer fMRI have been discussed elsewhere (Merriam & Kay, 2022; Viessmann & Polimeni, 2021). In our opinion, one important issue in the context of replicability and reproducibility is ROI selection. Especially in association cortices it is very difficult to localize an ROI due to less correspondence between macroscopic anatomy and function as well as unknown structure-function relationships at the mesoscale (Frost & Goebel, 2012). Given these uncertainties, the exact locations of specific functional regions are also much more variable between subjects than those in primary sensory and motor cortices (Mueller et al., 2013). In this context, identifying ROIs functionally within each subject individually using adequate localizer tasks (Fedorenko et al., 2024) or dense-sampling resting-state approaches, where resting-state networks can be precisely mapped on an individual-level basis (Gordon et al., 2017) can help address these challenges by reducing inter-subject variability. Lastly, variability in local cortical geometry and local vasculature may influence resulting laminar response profiles adding a further ROI selection dependence to the results. A better understanding of all these factors as well as objective, yet precise methods of ROI selection are needed.

In addition to determining ROI location and extent tangentially to the cortical surface, the manual ROI selection employed by Finn et al. (2019) also entailed gray matter segmentation and the delineation of superficial and deep layers. This second aspect, while not ideal for reproducibility, can potentially offer more precise results. One disadvantage is the significant effort required, which, although still manageable in this study, could become burdensome in larger studies or those involving multiple or larger ROIs, or targeting the entire cortex. Furthermore, we caution against assuming the absolute superiority of manual segmentations. While manual segmentation combined with ROI selection may lead to avoidance of regions affected by imaging artifacts or venous formations, our experience suggests that this also may introduce a bias towards geometrically simpler ROIs. For instance, in our manual ROI segmentations, we adhered to an automatically generated location and extent and observed that some geometrically complex ROIs might not have been selected manually, particularly when relying on predefined orthogonal cuts. The orientation of these cuts can also affect the accuracy of manual layer delineations. Lastly, manual segmentation may lead to false confidence in subjective interpretations of images affected by noise and artifacts, potentially overestimating its advantages compared to automatic segmentation results.

Lastly, given the wide range of analyses approaches currently used in layer fMRI and the above discussed issue of ROI selection, we believe it to be beneficial that the pioneering experiments are being followed up by studies in which hypotheses and analytic choices have been preregistered. The prevalence of feasibility studies in current layer fMRI research, that focus on confirmation of basic models of laminar involvement, renders them particularly conducive to pre-registration.

In summary, we were not able to fully reproduce the laminar dissociation originally reported by Finn et al. (2019). While we replicated the superficial-layer effect during WM manipulation, we found no evidence for deep-layer activation during motor response execution. This outcome is biologically plausible in light of invasive primate data and methodological considerations inherent to laminar fMRI. Electrophysiological studies reveal heterogeneous laminar profiles—some implicating deep layers, others superficial— depending on task demands and circuit context, and the fMRI signal itself primarily reflects synaptic input and dendritic integration rather than spiking output. Combined with task design factors that conflate retrieval and motor processes, a clean, exclusive deep-layer effect is unlikely. Our findings therefore refine current models of laminar organization in human dlPFC, consolidating superficial-layer engagement as a reproducible signature of WM manipulation and aligning human evidence with the more nuanced animal literature. By emphasizing preregistration, automation, and transparent analytic choices, this study sets a benchmark for reproducibility in laminar fMRI and strengthens the methodological and conceptual foundation needed to link systems-level cognition with microcircuit-level mechanisms.

## Supporting information

Supplementary Results and Figures

## Acknowledgements

We thank Emily Finn and Renzo Huber for helpful discussions and for providing additional information about analyses related to Finn et al. (2019). J.K.D. was supported by the Max Planck Society and BMBF. N.W. was supported by the Deutsche Forschungsgemeinschaft (DFG, German Research Foundation) – project no. 347592254 (WE 5046/4-2); the Federal Ministry of Education and Research (BMBF) under support code 01ED2210; the European Union’s Horizon 2020 research and innovation programme under the grant agreement No 681094. R.L. was funded by the Wellcome Trust (209139/Z/17/Z). This project also received funding from the Klaus Tschira Stiftung gGmbH foundation (support code GSO/KT18). This paper was typeset with the bioRxiv word template by @Chrelli: www.github.com/chrelli/bioRxiv-word-template

## Data and code availability

All code is accessible under https://github.com/Cognitive-Neuroscience-Neurotechnology/2025_pfc-layer-wm-replication_7tmri_dchaimow. All data, code, and analysis software environments will be uploaded to publicly accessible repositories upon publication.

## Methods

### Participants and sample size rationale

22 healthy participants were recruited from the Max Planck Institute for Human Cognitive and Brain Sciences (MPI CBS) subject database. All participants had normal or corrected to normal vision, were native German speakers, and right-handed as determined by the Edinburgh handedness inventory. Participants had no history of neuropsychiatric disorders and were not currently taking psychoactive medications. All participants gave written and informed consent. The study was approved by the ethics committee of the University of Leipzig (441/20-ek).

Data from one subject was excluded from further analysis due to excessive head motion as prescribed in the preregistration (relative motion in at least two runs, as estimated by retrospective motion correction, exceeded the mean plus 3 standard deviations from all runs in all subjects).

The final sample consisted of 21 participants (ages 18 – 36 years, 10 female), which was our predefined target sample size and had been determined as follows. First, we performed a power analysis in G*Power (Faul et al., 2009), using partial eta squared values calculated from the ANOVA results reported in Finn et al. (2019) (personal communication). Specifically, we took the smallest interaction effect of the below described across layers analysis (*η*_p_ ^2^ = 0.681 for the interaction between layer and contrast during the response period). We decreased this effect size by ⅓ to 0.45 to account for a fewer planned number of runs per subject in our study. Due to time constraints imposed by the maximum allowable duration for scanning per subject in one session (90 min), we limited the data acquisition to 4 instead of 5-6 runs; this decision was made in accordance with local institutional regulations and ethical guidelines specified for our study. We set the effect size specification option in G*Power to “as in Cohen (1988)”. Note that this gave us the most conservative results. For the non-sphericity correction, we used the smallest sphericity index estimate from the across layers analysis (ε=0.612; communicated to us by the authors). We further set α =0.05, the desired power to 1-β=0.8, the number of groups to 1 and the number of measurements to 4. This approach resulted in a required sample size of 21 subjects. We performed a bootstrapping simulation using all single subject results from Finn et al. (2019) provided to us by the authors. From that data we randomly generated 1000 sets of single subject datasets for each tested subject sample size (sampling with replacement). We then ran the same across layers ANOVA as in Finn et al. (2019) and counted the proportion of statistically significant interaction effects (p<0.05). This procedure resulted in a minimum sample size of only 6 subjects to achieve a power of at least 80%. Note that in contrast to our G*Power analysis, the bootstrapping analysis did not consider the reduced amount of data per subject. Nonetheless, the bootstrapping analysis indicates the strength of effects reported in Finn et al. (2019). However, to be conservative we chose the larger sample size of 21 subjects, as estimated above, as our target sample size.

### Experimental procedure

Participants performed a working memory task inside the scanner. The task was written in GNU Octave, using the Psychophysics Toolbox extensions (Brainard, 1997; Kleiner et al., 2007; Pelli, 1997) and was made identical to the description in Finn et al. (2019) (see their “axial protocol”). Identical to Finn et al. (2019) there were two types of experimental runs (alphabetization vs. remembering and action vs. non-action), each consisting of 20 pseudo randomized trials. After instruction and a short behavioral training to familiarize with the task, participants underwent a 6-min whole-brain functional localizer run using a lower resolution GE-BOLD sequence. Identical to Finn et al. (2019) this localizer consisted entirely of “ALPHABETIZE” trials and had slightly altered timing (the ITI was shortened from 16 s to 5 s). The functional localizer scan was then analyzed for delay period activity, and a high-resolution SS-SI-VASO slab was positioned to maximize coverage of activated regions within and around the left dlPFC. After the acquisition of structural T1 weighted images participants performed up to two runs of each type in an interleaved manner for a total of 4 runs. Finn et al. (2019) used up to 6 runs per subject, a number which we could not achieve, due to maximum scanning time constraints at the MPI CBS. Note that this was accounted for in our target sample size calculation.

### MRI protocol

MRI data was acquired using a Siemens MAGNETOM Terra 7T MRI pTx system (Siemens Healthineers, Erlangen, Germany) at the MPI CBS, using an 8Tx/32Rx channel head coil (Nova Medical Inc., Wilmington, MA, USA). Structural T1 weighted images were acquired using MP2RAGE sequence (Marques et al., 2010) (TR = 5000 ms, TE = 2.27 ms, 0.75 mm isotropic voxels) at two inversion times (TI of 900 ms, 2700 ms with a flip angle of 3°, 5°, respectively) that were combined to yield a T1-weighted image with uniform contrast (UNI). High-resolution partial-brain functional data were acquired using an SS-SI-VASO sequence with 3D-EPI readout (Huber et al., 2020; Stirnberg & Stöcker, 2021) (sequence version number 822d59f4). The resolution was 0.8 x 0.8 x 0.9 mm^3^, with a FOV of 150 x 150 x 21.6 mm^3^. The volume TR encompassing the acquisition of a nulled and not-nulled image was 3.7 s in total. Further parameters were TE=18.7 ms, TI1=1.4 s, TI2=2.9 s, GRAPPA factor 3, 6/8 partial Fourier in the first phase encoding direction, readout bandwidth=1064 Hz/pixel). A lower resolution whole-brain 2D EPI scan (2 mm isotropic resolution, TR = 2 s, TE = 18 ms) was used for intra-session functional localization and slab positioning.

### Differences in data acquisition to Finn et al. 2019

Our MRI scanner (Siemens 7T Magnetom Terra) differed from the one used in Finn et al. (2019). Although both scanners were produced by Siemens Healthineers, they ran different non-compatible software versions. Therefore, we were not able to use the exact SS-SI-VASO pulse sequence that was used in Finn et al. (2019). Instead, we used a newer, slightly different implementation (Huber et al., 2020; Stirnberg & Stöcker, 2021). Among other things, this sequence differed in the implementation of the 3D-EPI read-out (Stirnberg & Stöcker, 2021), in that the temporal spacing of nulled and not-nulled read-out blocks is not symmetric and that it did not use a phase-skip parameter. Consequently, possible and recommended parameter protocols differed somewhat between both sequences. Our employed resolution was 0.8 x 0.8 x 0.9 mm^3^, compared to 0.9 x 0.9 x 1.1 mm^3^ in Finn et al. (2019), which is expected to yield more thermal noise but also better resolvability between layers. Our TR, including the measurement of 1 nulled and 1 not-nulled volume, was 3.7 s compared to 4 s in Finn et al. (2019).

### Data conversion and anonymization

Before further processing all data was converted from DICOM to Nifti format, anonymized, and structural data was defaced (Gulban et al., 2019). In subject 1, the high-resolution functional runs consisted of 188 instead of 178 volumes. These additional volumes were discarded. All data was organized into a BIDS compatible structure, adding meta information as necessary (e.g. task event files).

### Preprocessing of structural data

We developed a custom processing pipeline around Freesurfer’s recon-all (Fischl, 2012). This addressed the problem that residual noise in processed MP2RAGE data results in difficulties with skull stripping and that high-resolution 7T T1w images are difficult for estimating gray matter without being affected by dura mater. Using pilot data sets, we informally explored several different approaches before settling on what produced the most accurate gray matter and white matter surfaces. The MP2RAGE INV2 images (second inversion, mostly proton-density weighted) were bias-corrected using SPM12 (Ashburner & Friston, 2005) before multiplying it with the MP2RAGE UNI images (uniform T1-weighted images with salt-and-pepper noise in regions of low signal) to yield MPRAGE like images (Kashyap, 2021). These images were fed into the first stage (autorecon1) of Freesurfer’s (v7.3.2) recon-all processing stream without skull-stripping using the highres flag. The same images were fed into the CAT12 (Gaser et al., 2024) segmentation stream, and a brain mask was defined as the union of resulting gray and white matter segmentations. This brain mask was then used to substitute Freesurfer’s skull stripping steps, specifically applying it to orig.mgz, writing the result as brainmask.mgz and brainmask.auto.mgz. After that, the remaining stages (autorecon2 and autorecon3) of Freesurfers recon-all processing stream were run, using the highres flag and setting the mris_inflate iterations to 100 using an expert option file.

For subsequent use of the surface-based HCP MMP 1.0 atlas (Glasser et al., 2016) within each individual subject, the structural surfaces of each participant were brought into fsLR 164k space using ciftify_recon_all, which is part of the Python-based ciftify package (Dickie et al., 2019), with expert settings to register to the high-resolution 0.5 mm MNI152 T1 template. Ciftify adapts the post-Freesurfer portion of the HCP minimal preprocessing pipeline (Glasser et al., 2013) to non-HCP acquired data and includes surface-based alignment (Robinson et al., 2014) to HCPs fs_LR space.

### Preprocessing of functional data

The DICOM images of each run were converted into a nulled and not nulled series. The first two volumes of each series were discarded to account for initial T1 relaxation effects. Motion correction across volumes and runs was performed using AFNI’s 3dvolreg and was done separately for the nulled and not-nulled series. Volumes from the same run and with the same index were used as references for both and were determined as those that together had the lowest number of outliers in their voxels as determined by AFNI’s 3dToucount. Finally, runs belonging to each type (alphabetization vs. remembering, action vs. non-action) were averaged.

### Trial averaging

Different from Finn et al. (2019) the trial duration, which we kept identical to 32 s, was not a multiple of our TR (3.7 s). Therefore, we used finite impulse response modeling to estimate the average trial time course. We estimated raw signal trial averages separately for the nulled and not-nulled data using AFNIs 3dDeconvolve with the TENTzero model and a separate set of polynomial regressor up to order 5 to account for low temporal frequency drift. For the not-nulled time series, we shifted the model in time to account for the delay between nulled and not-nulled acquisition (1.5 s), so that the resulting trial averages were temporally aligned with each other. The nulled trial average was then divided by the not-nulled trial average resulting in a BOLD corrected VASO estimate while the not-nulled trial average was from here on considered as the corresponding BOLD estimate. Finally, the VASO and BOLD trial averages for each voxel were transformed to units of percent signal change by dividing by the constant predictor value (BOLD corrected in the case of VASO). Note that the VASO signal is expected to be inversely proportional to cerebral blood volume, therefore all VASO plots depict negative percent change which is interpreted as a positive activation. In addition, this analysis yielded voxel-wise F-statistics that quantified the amount of variance explained by the trial structure of the experiment. The F-statistics from the not-nulled time series were subsequently used as a BOLD based measure of task involvement.

### SNR calculation

In order to calculate a representative VASO tSNR measure we additionally performed BOLD correction on data from individual not-averaged runs. The not-nulled time series was temporally realigned with nulled time series using FSL’s slicetimer and nulled volumes were divided by their not-nulled counterpart yielding BOLD corrected VASO time series for each run. Time-course SNR (tSNR) was calculated by dividing the mean by the standard deviation of each voxel’s motion- and BOLD-corrected VASO time series in each individual run. tSNR values were then first averaged across runs and finally across all voxels within each subject’s ROI.

### Co-registration and calculation of cortical depths

A T1w image was calculated from all nulled and not-nulled VASO images as follows. All runs were averaged and the inverse coefficient of variation (mean divided by variance) over all volumes (nulled and not-nulled) was computed. This procedure emphasizes the T1 contrast originating from different T1 weighting of nulled and not-nulled read-outs and makes it qualitatively similar to the processed MP2RAGE images. The resulting images were further processed using a custom pipeline that included background denoising using LAYNII (Huber et al. 2021) (LN2_MP2RAGE_DNOISE) and bias field correction (SPM12). Next, the Freesurfer processed T1w image (from MP2RAGE acquisition) was registered to the VASO T1w image using ANTs (Tustison et al., 2021). The registration consisted of 3 stages: rigid, followed by affine and finally non-linear SyN registration. The rigid and affine stages used mutual information as metric and the SyN stage (Avants et al., 2008) used cross-correlation. The resulting transformation was then applied to the Freesurfer gray and white matter surface coordinates, aligning the surfaces with the functional data. From these surfaces, voxel-wise cortical depths were computed using a custom algorithm. In effect, this algorithm calculated for every voxel the relative depth at which a transformation between gray and white matter boundary (under an equivolume model) would intersect with the voxel’s center (Chaimow et al., 2022; https://github.com/dchaimow/vdfs). Superficial and deep layers were defined as voxels having a relative cortical depth of <0.5 and >0.5 respectively. In a control analysis that introduced a gap between layers these depth ranges were changed to <⅓ and >⅔. HCP MMP 1.0 atlas labels (Glasser et al., 2016) were transformed from fs_LR to each subject’s native Freesurfer surface space using HCP’s wb_command (label-resample with ADAP_BARY_AREA method) and the subjects surface transformations previously generated by cifify’s recon-all stream. The labels were then projected into functional volume space using HCP’s wb_command (label-to-volume-mapping with ribbon-constrained option) using the individual subjects’ surfaces in functional volume space.

### Definition of ROI

For ROI definition we combined atlas information with functional activation results. First, the BOLD trial averaging F-statistics from alphabetization vs. remembering and action vs. no action runs were averaged, sampled onto the cortical surface (wb_command -volume-to-surface-mapping with ribbon-constrained option) and smoothed (wb_command -metric-smoothing with FWHM = 3 mm).

This map was masked so that only activity within left p9-46v and all surrounding parcels remained. Given a specific threshold value (that was computationally obtained - see description below), the surface ROI was defined as the largest connected cluster that showed any overlap with the center parcel (left p9-46v).

Using an optimization algorithm (scipy brentq; Brent, 1973), we then estimated the threshold value that resulted in an ROI whose surface area (calculated with wb_command metric-weighted-stats) was closest to a specified target surface area. For the main analyses, this target surface area was chosen to be 100 mm^2^ after qualitative comparison of the size of ROIs for different target surface areas with the size of ROIs presented in Supplementary Fig. 3 of Finn et al. (2019). Note that we were not able to quantify this as ROIs were not published in the original study. The final surface ROI was then projected into the corresponding gray matter of the EPI volume (using wb_command metric-to-volume-mapping with the ribbon-constrained option). ROIs were split into a superficial and deep layer by binning the voxels according to their relative depth.

### Definition of alternative ROIs constrained by group analysis clusters

The individual subjects’ BOLD trial averaging F-statistics, trial type combined and surface sampled, were transformed to fs_LR space, averaged across subjects and smoothed (sigma=3). The group averaged F-statistics map was then used to define clusters using a custom algorithm. The algorithm iterates over vertices sorted by their values in descending order (until a minimum average F-value of 1 is reached), tracking and merging clusters of connected vertices as they become adjacent. Clusters that have reached a specified minimum size (100 mm^2^) before they would merge with other neighboring clusters are considered final. This procedure ensures that each cluster represents a locally maximal region. Group clusters were then back-projected into single subject surfaces and individually used as constraining regions for the procedure described in ‘Definition of ROI’.

### Control analysis using manually segmented laminar ROIs

ROIs consisting of a superficial and a deep layer were drawn manually on ROIs consisting of a superficial and a deep layer were drawn manually on top of the T1 weighted VASO image. For that, we followed the description provided by Finn et al. (2019). The previously computed automatic ROIs were used to determine the location and extent, but gray matter segmentation and layer assignment was purely based on the T1 weighted VASO image. The T1 weighted MP2RAGE image was occasionally only consulted to provide additional contextual information about the local cortical geometry.

### Statistical analysis of trial period responses

Statistical analyses followed Finn et al. (2019). Trial averaged percent change time courses from all voxels belonging to the ROI were averaged according to cortical depth (deep vs. superficial) and condition. Contrast time courses were calculated as the alphabetization vs. remembering and action vs. non-action difference. Trial period responses were extracted by averaging time points 3, 4 (0-based) for the delay period (taking hemodynamic delay into account) and 5, 6 for the response period following the probe. Identical to Finn et al. (2019) we compared differential activity within trial periods and subsequently non-differential activity within layers using a series of two-way (2 x 2) repeated measures ANOVAs, testing for interaction effects using a threshold of p<0.05. To answer how the difference of activity between trial types depends on the layer and the contrast of interest:

1. For the delay period:

factor 1: layer (superficial, deep)
factor 2: contrast (alphabetization vs. remembering, action vs. non-action)
Dependent variable: difference of activity during delay period according to contrast
2. For the response period:

factor 1: layer (superficial, deep)
factor 2: contrast (alphabetization vs. remembering, action vs. non-action)
Dependent variable: difference of activity during response period according to contrast

We consider this set of ANOVAs (1. and 2.) to be our primary analyses of interest. These are the analyses that we have optimized our target sample size for.

To answer how the activity within layers depends on trial type and trial period:

1. For superficial layers and alphabetization vs. remembering runs:

factor 1: trial type (alphabetization, remembering)
factor 2: trial period (delay, response)
dependent variable: activity during trial period
2. For superficial layers and action vs. non-action runs:

factor 1: trial type (action, non-action)
factor 2: trial period (delay, response)
dependent variable: activity during trial period
3. For deep layers and alphabetization vs. remembering runs:

factor 1: trial type (alphabetization, remembering)
factor 2: trial period (delay, response)
dependent variable: activity during trial period
4. For deep layers and action vs. non-action runs:

factor 1: trial type (action, non-action)
factor 2: trial period (delay, response)
dependent variable: activity during trial period

### Confidence interval bootstrapping analysis

1000 bootstrap samples of size 21 each were generated by randomly selecting subjects with replacement. For each sample an ANOVA analysis was run as described above (factors: layer and condition contrast). In addition, the direction of the mean layer difference (superficial>deep or deep>superficial) with respect to the alphabetization vs. remembering contrast (in case of delay period) or with respect to the action vs. non-action contrast (in case of response period) was calculated. This resulted in distributions of ANOVA interaction effect sizes (η_p_^2^) for each trial period. Confidence intervals to assess overlap with effect sizes from Finn et al. (2019) were calculated by calculating 95th percentiles, first considering all samples associated with a layer difference opposite to that of Finn et al. (2019) in decreasing order, followed by all samples associated with the same layer difference in increasing order.

