## Supplementary Results and Figures for "Re-evaluating Laminar Specificity of Working Memory in Human Prefrontal Cortex"

bioRxiv

[a]Cognitive Neuroscience & Neurotechnology Group, Max Planck Institute for Biological Cybernetics, Tübingen, Germany
[b]Department of Neurophysics, Max Planck Institute for Human Cognitive and Brain Sciences, Leipzig, Germany
[c]Department of Radiology and Medical Informatics, University of Geneva, Switzerland
[d]Neuro-X Institute, Ecole Polytechnique Fédérale de Lausanne, Switzerland
[e]Athinoula A. Martinos Center for Biomedical Imaging, Massachusetts General Hospital, Charlestown, MA, USA
[f]Department of Radiology, Harvard Medical School, Boston, MA, USA
[g]Felix Bloch Institute for Solid State Physics, Faculty of Physics and Earth System Sciences, Leipzig University, Leipzig, Germany
[h]Wellcome Centre for Human Neuroimaging, Institute of Neurology, University College London, 12 Queen Square, London WC1N 3AR, UK

### Supplementary Result 1: Manipulation-dependent signal increases in both layers

Activity in both layers was higher in alphabetization trials than in remembering trials independent of the trial period (superficial layers: $p=8.6\times10^{-5}$, $F(1,20)=24.0$, $\eta_p^2=0.546$; deep layers: $p=3.6\times10^{-3}$, $F(1,20)=10.9$, $\eta_p^2=0.352$; effect of condition in a two-way ANOVA with factors condition x trial period). We found no significant interactions between trial type and period (superficial layers: $p=0.61$; $F(1,20)=2.6\times10^{-1}$; $\eta_p^2=1.30\times10^{-2}$;deep layers: $p=0.83$; $F(1,20)=4.7\times10^{-2}$; $\eta_p^2=2.35\times10^{-3}$).

### Supplementary Result 2: Action dependent signal increase in superficial but not deep layers

For the action vs. non-action comparison, we found an interaction between trial type and trial period in superficial layers ($p=9.9\times10^{-4}$; $F(1,20)=14.8$; $\eta_p^2=0.426$), such that activity in action trials was higher than in non-action trials during the response period. Also in deep layers, we found an interaction between trial type and trial period. Again, activity was higher in action trials than in non-action trials ($p=8.9\times10^{-3}$; $F(1,20)=8.3$; $\eta_p^2=0.296$).

### Supplementary Figure 1: Main results using BOLD data

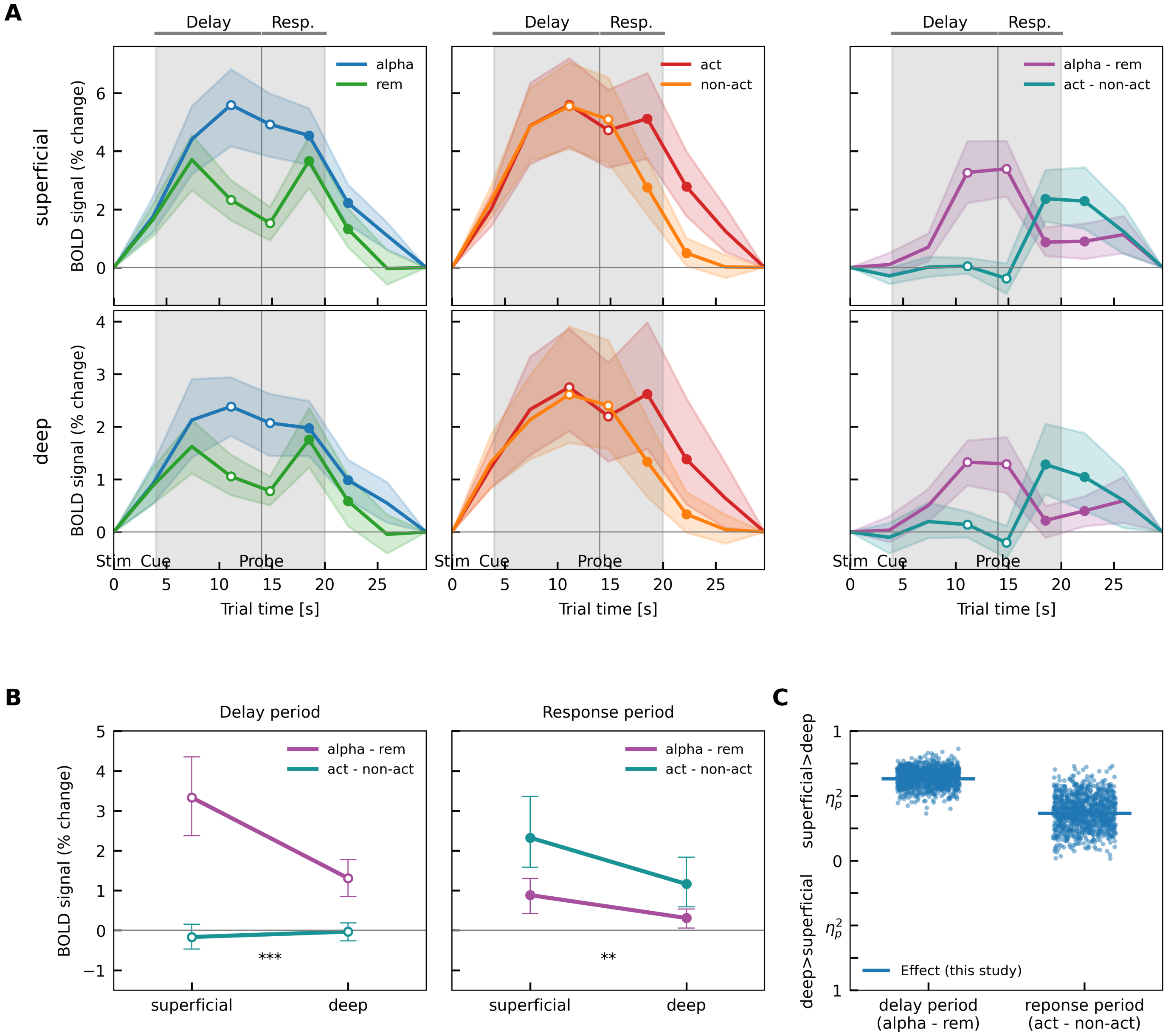


**Supplementary Figure 1: Main results using BOLD data**. Same format as Fig. 1.

### Supplementary Figure 2: Time-course SNR (tSNR)

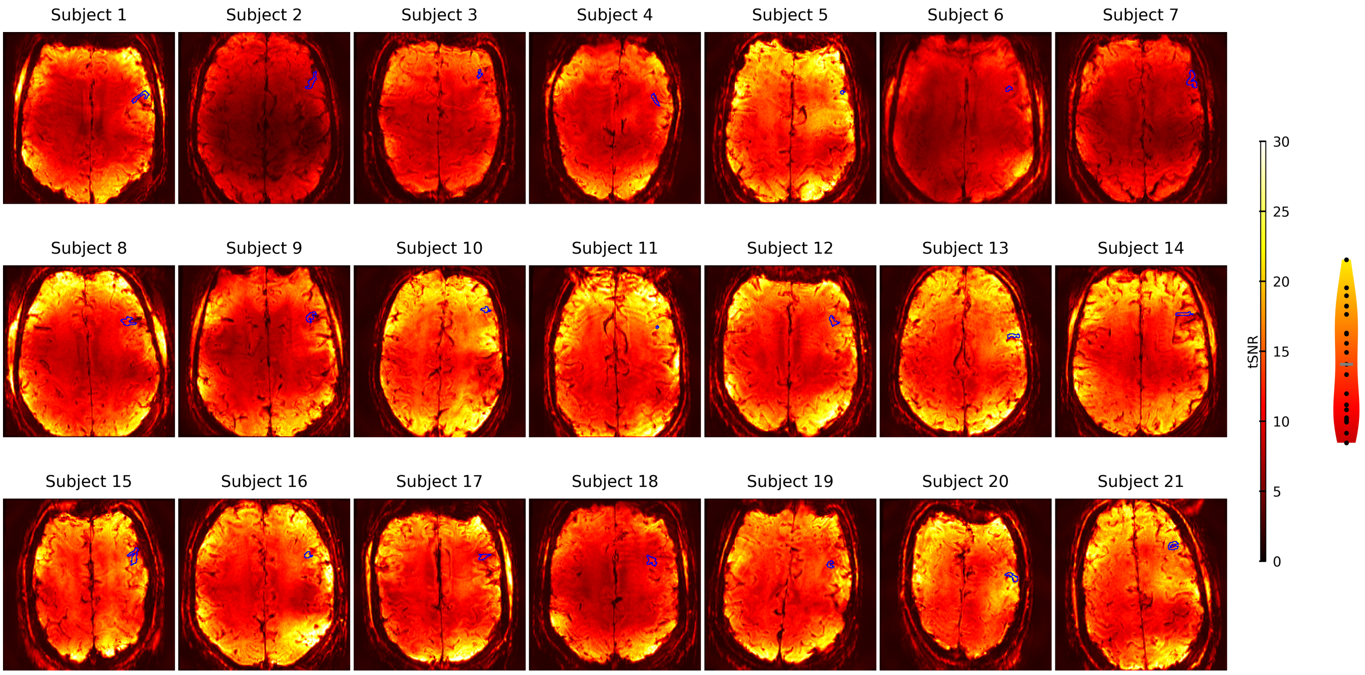


**Supplementary Figure 2:** Time-course SNR (tSNR). For each subject the tSNR map calculated from individual runs and averaged over all runs is shown from a single slice cutting through the ROI. The outline of the ROI is superimposed in blue. The plot on the right shows the distribution across subjects of tSNR values averaged over each ROI (median = 14.1, in gray).

### Supplementary Figure 3: Segmentation, Registration and individual ROIs

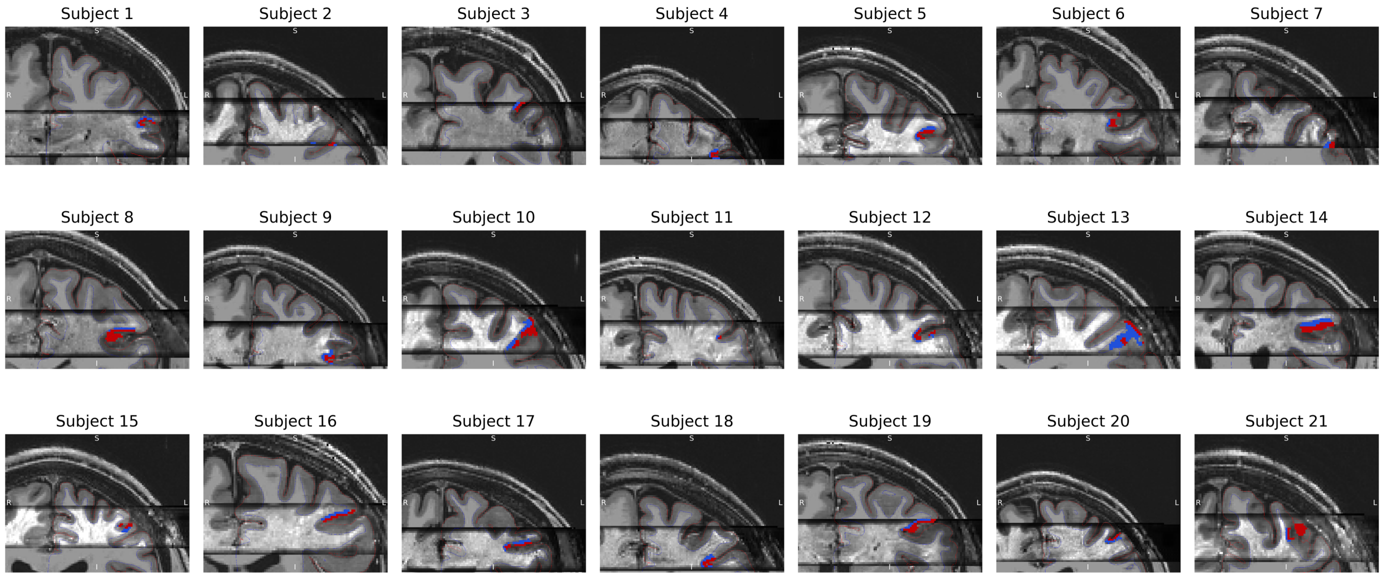


**Supplementary Figure 3: Segmentation, Registration and individual ROIs.** For each subject, the T1w-like image calculated from the VASO data (slab) is shown superimposed on the registered T1w image obtained from MP2RAGE. Reconstructed white matter and pial surfaces were transformed to the slab space using the same registration and are shown in red and blue, respectively. Superimposed, each subject's ROI is shown separated into superficial (red) and deep layers (blue).

### Supplementary Figure 4: Main results without slab boundary

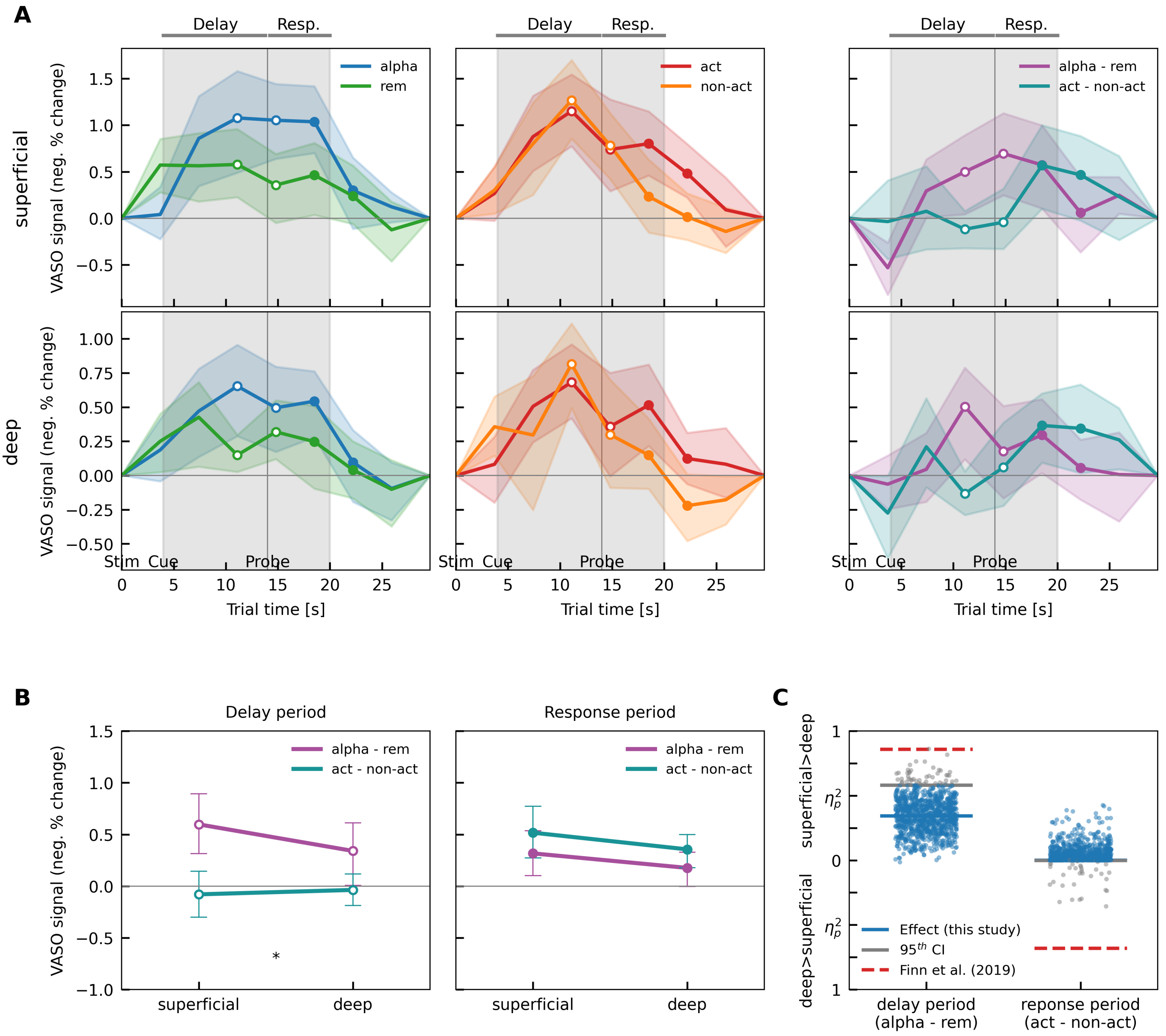


**Supplementary Figure 4: Main results from a subset of subjects (N=17) whose ROIs were not close to the slab boundary.** Same format as Fig. 1.

### Supplementary Figure 5: Results from manually defined laminar ROIs

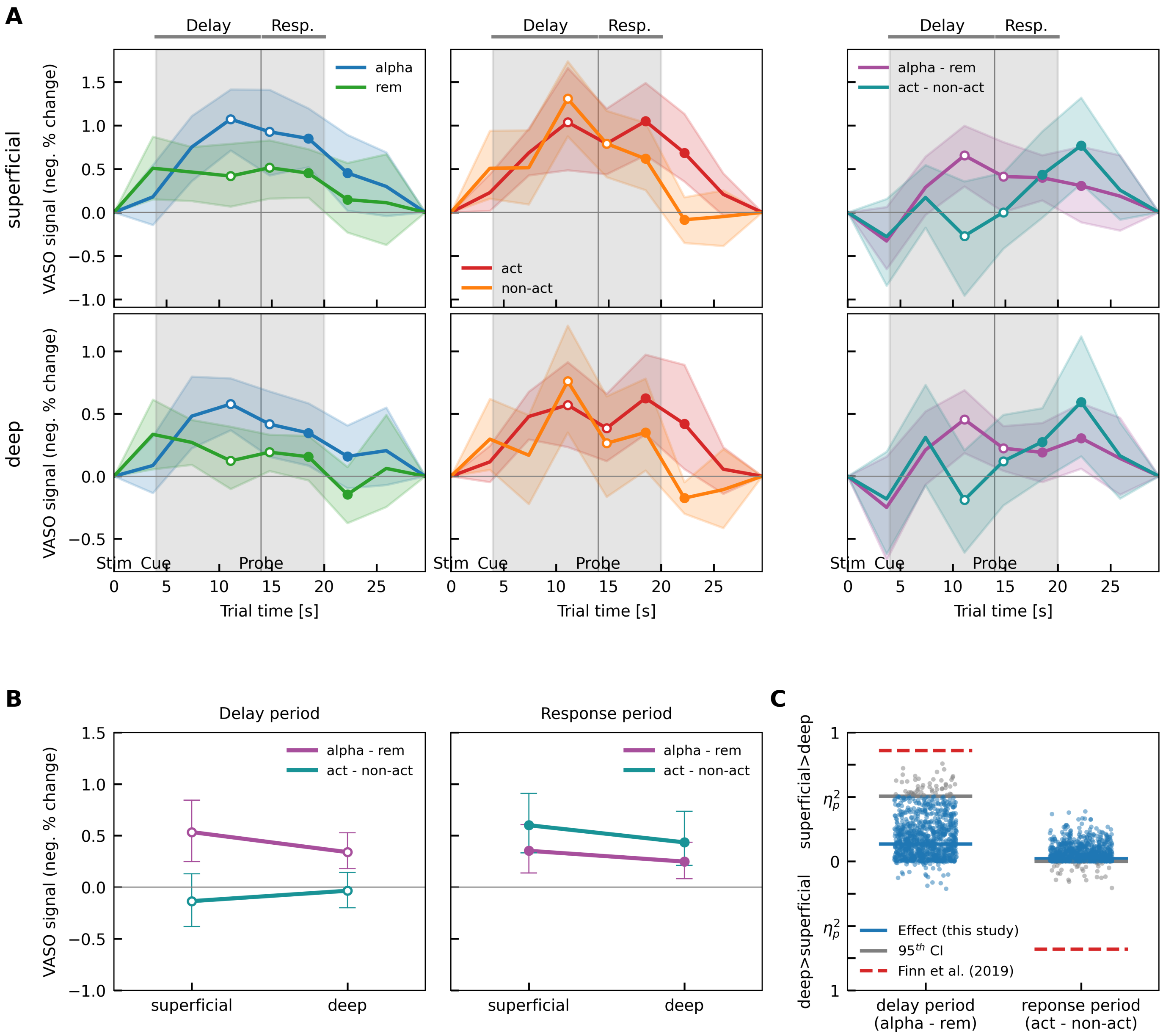


**Supplementary Figure 5: Main results using manually defined laminar ROIs.** Same format as Fig. 1.

### Supplementary Figure 6: ROI size dependence of results

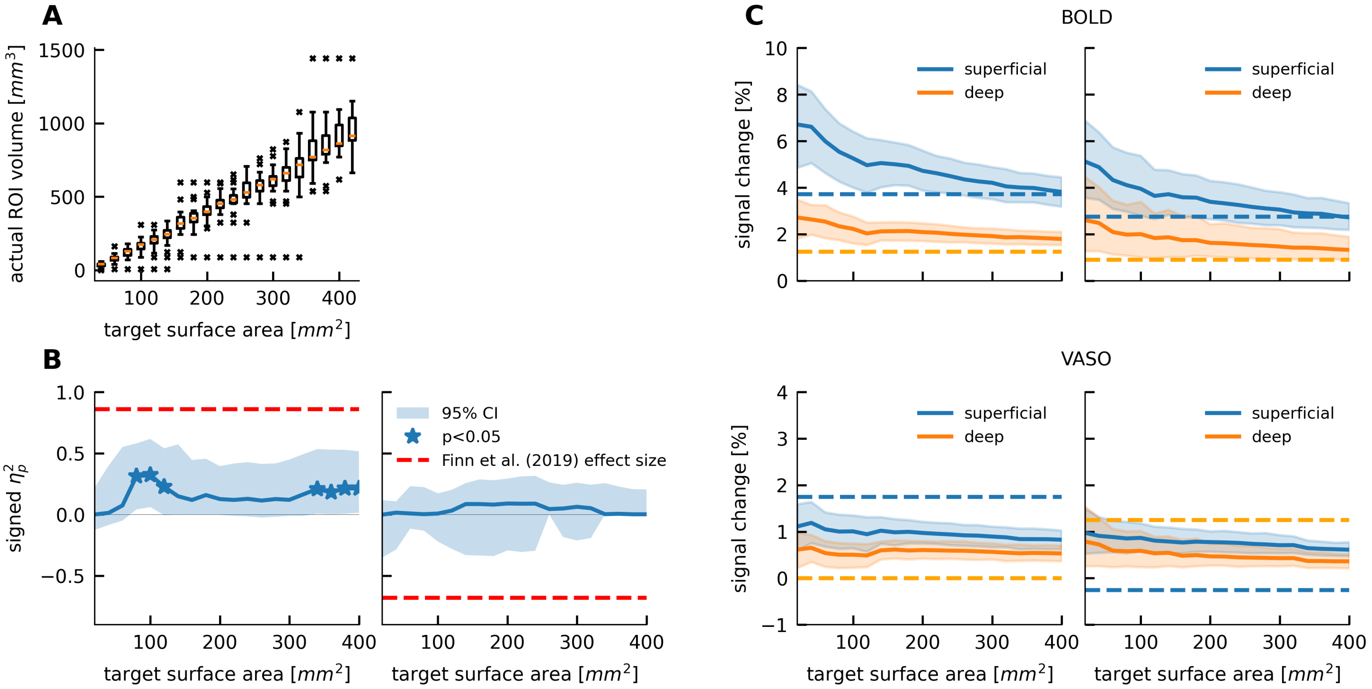


**Supplementary Figure 6: ROI size dependence of results. A** shows the distribution of actual ROI volumes as a function of the target ROI surface area that was given to the ROI estimation algorithm. In **B** the dependence of VASO layer effects (ANOVA interaction effect) for the delay period (alphabetization >remembering contrast; left) and response period (action > non-action contrast; right) is shown as a function of target ROI surface area. The effect size from Finn et al. (2019) is shown in red. **C** shows the dependence of activation amplitudes on ROI size for VASO (top) and BOLD (bottom) for the delay period (left) and response period (right). The dashed lines indicate the responses from Finn et al. (2019) (VASO from their Fig. 2, BOLD from personal communication).

### Supplementary Figure 7: Group analysis of activated regions in dlPFC

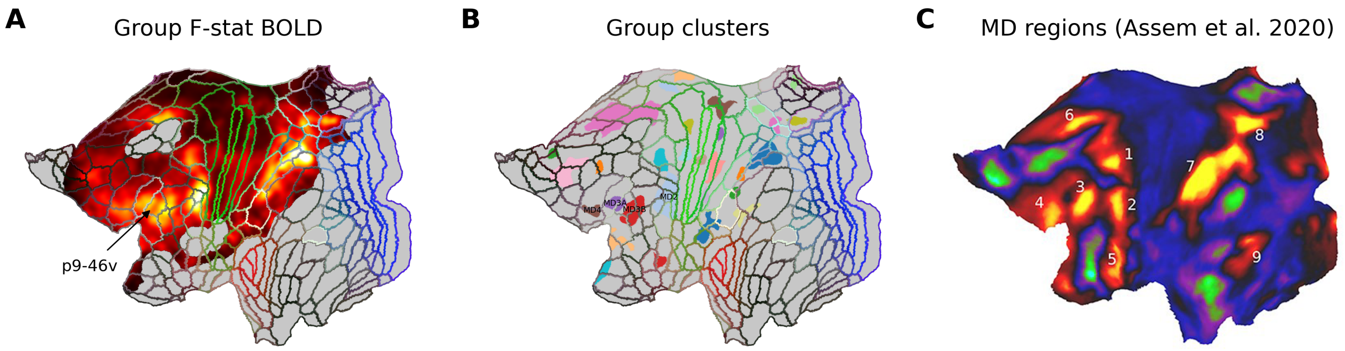


**Supplementary Figure 7: Group analysis of activated regions in dlPFC.** In order to explore alternative ROIs, we took the smoothed and group averaged BOLD F-stat map (**A**) and extracted individual clusters (**B**), that indicate high involvement with any component of the task. Many of these clusters show a strong resemblance to the multi-demand system regions (**C**; from Assem et al., 2020). We then reran our main analysis on VASO where we back-projected selected group clusters into individual subject space. We used these as location constraints for our ROI calculation algorithm that looked for local single subject BOLD activations. Results for these ROIs are shown in Supplementary Figures 8-11.

### Supplementary Figure 9: Results for neighboring dlPFC ROI based on group cluster MD3A

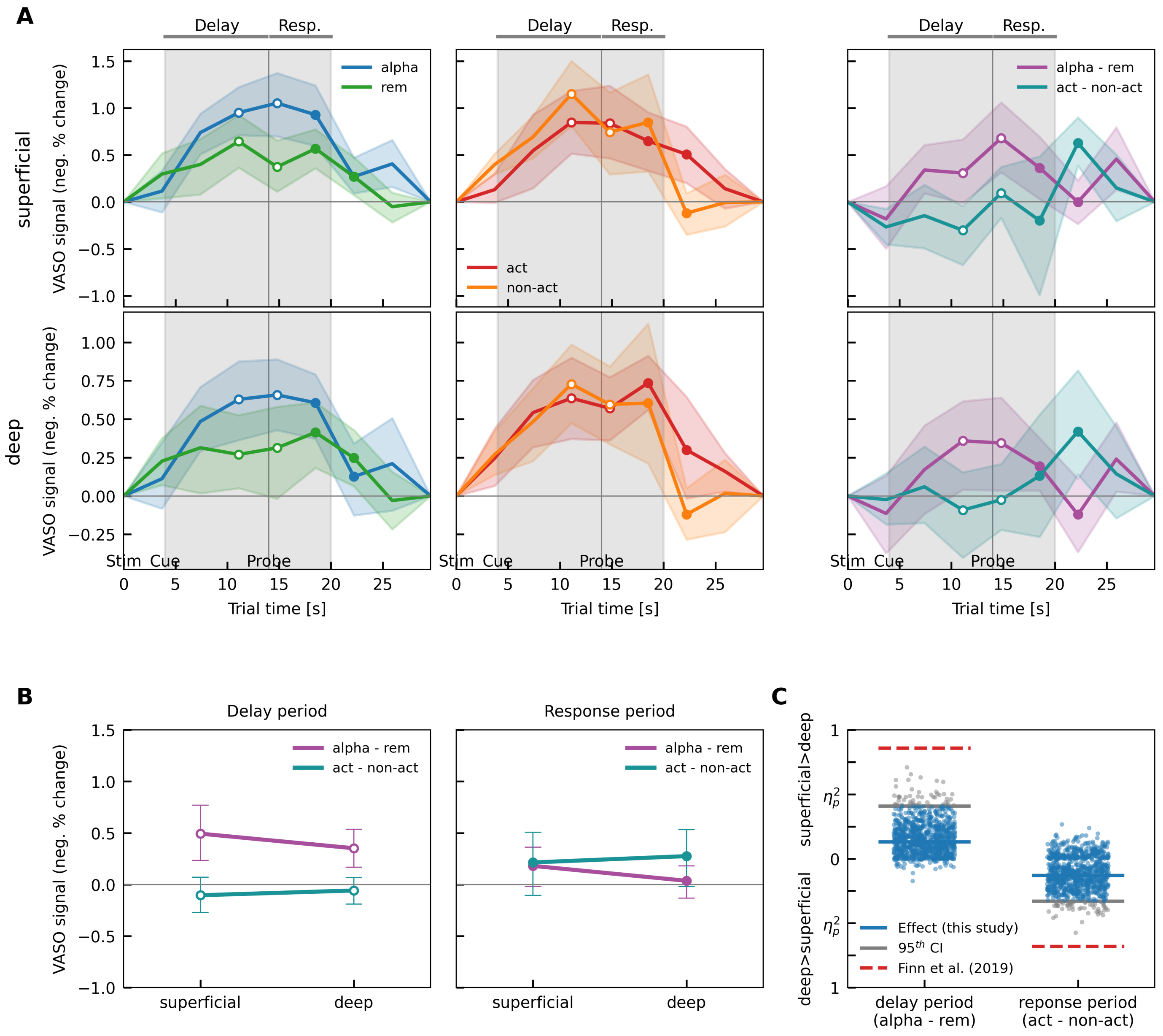


**Supplementary Figure 9: Results for neighboring dlPFC ROI based on group cluster MD3A.** Same format as Fig. 1.

### Supplementary Figure 10: Results for neighboring dlPFC ROI based on group cluster MD3B

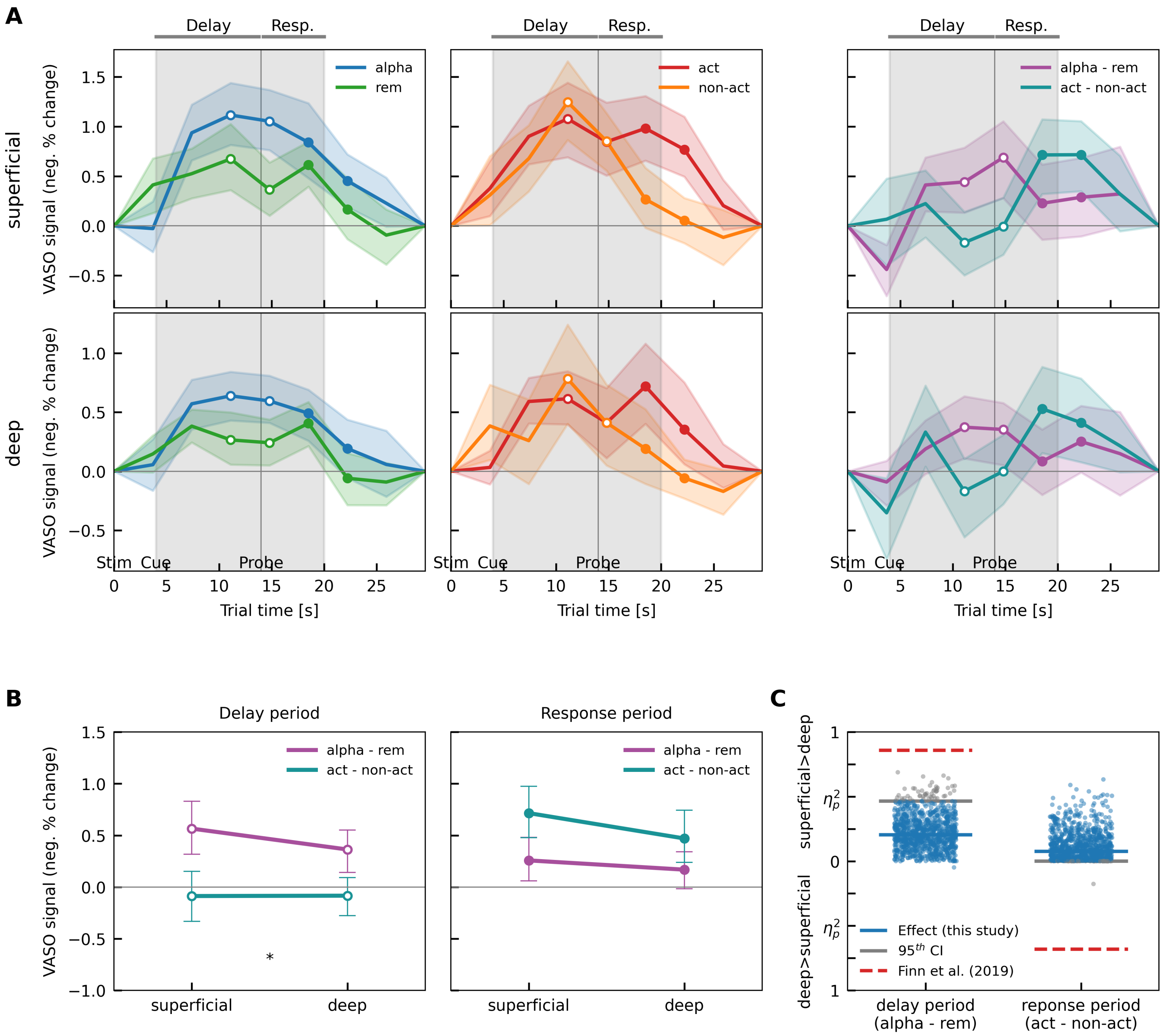


**Supplementary Figure 10: Results for neighboring dlPFC ROI based on group cluster MD3B.** Same format as Fig. 1.

### Supplementary Figure 11: Results for neighboring dlPFC ROI based on group cluster MD4

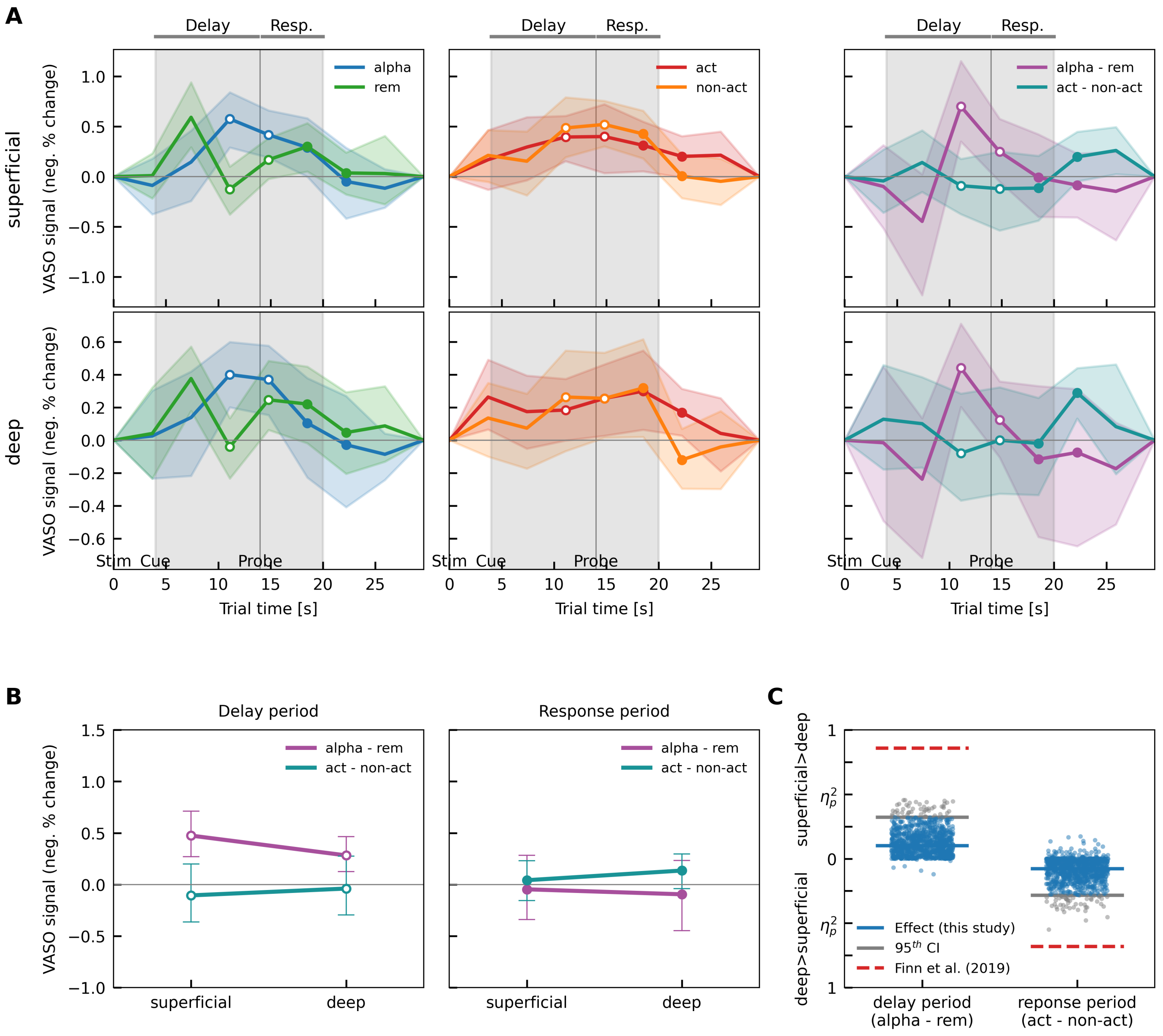


**Supplementary Figure 11: Results for neighboring dlPFC ROI based on group cluster MD4.** Same format as Fig. 1.

### Supplementary Figure 12: Results using between-layer gap

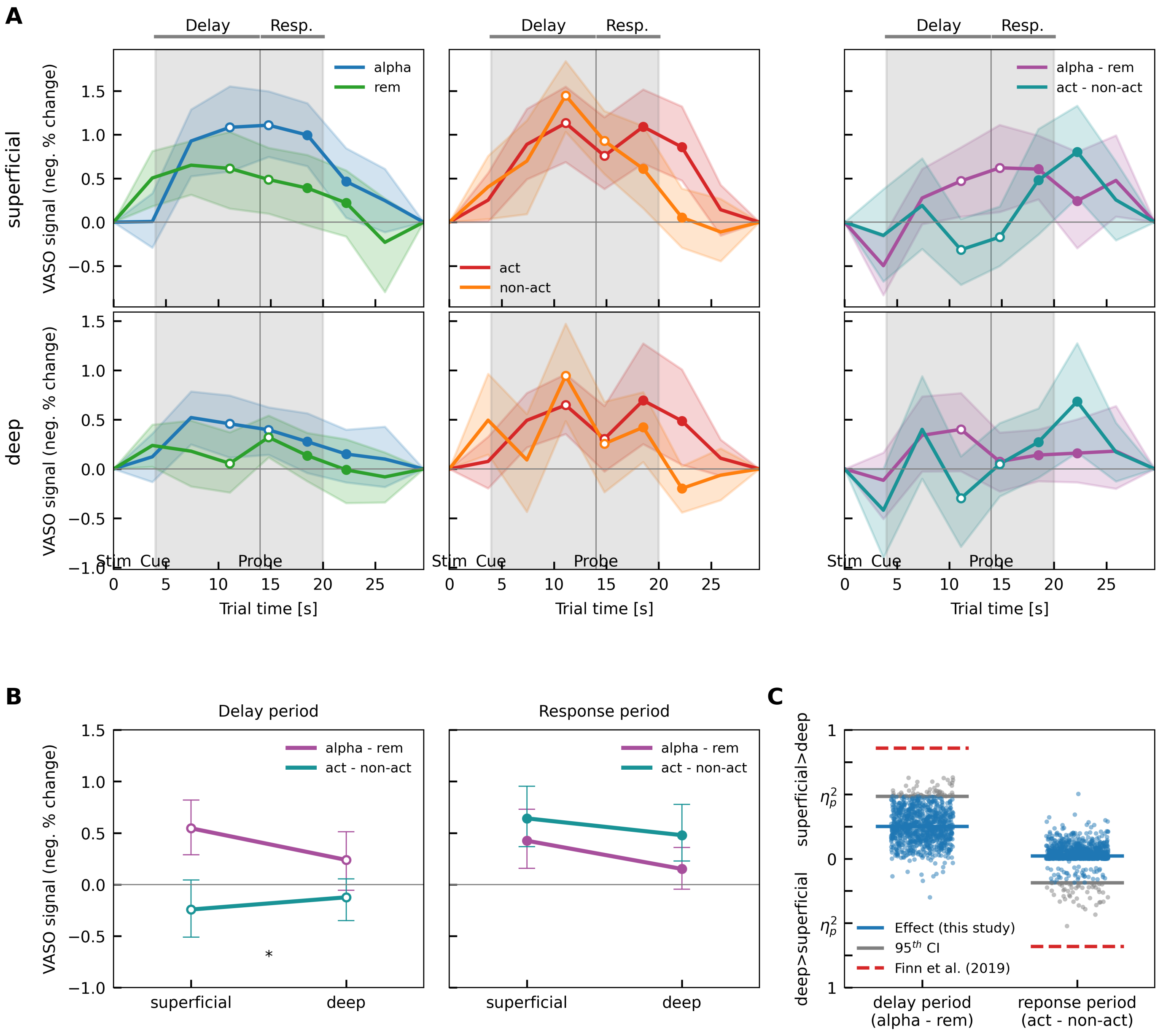


**Supplementary Figure 12: Main results when using a gap of ⅓ relative depth between deep and superficial layer ROIs.** Same format as Fig. 1.
